# Exploring human rare disease variants from a multidimensional perspective illuminates receptor – G protein coupling diversity

**DOI:** 10.1101/2024.01.16.575841

**Authors:** Theo Redfern-Nichols, Shannon L O’Brien, Xianglin Huang, Brian Medel-Lacruz, Davide Calebiro, Jana Selent, Graham Ladds, Maria Marti-Solano

**Affiliations:** Department of Pharmacology, University of Cambridge, Tennis Court Road, Cambridge CB2 1PD, UK; Institute of Metabolism and Systems Research, College of Medical and Dental Sciences, University of Birmingham, Birmingham B15 2TT, UK; Centre of Membrane Proteins and Receptors (COMPARE), Universities of Nottingham and Birmingham, Birmingham B15 2TT, UK; Research Program on Biomedical Informatics, Hospital del Mar Medical Research Institute, Department of Experimental and Health Sciences, Pompeu Fabra University, Barcelona, 08003, Spain

## Abstract

G protein-coupled receptors (GPCRs) are transmembrane proteins capable of detecting signals as diverse as odours, neurotransmitters, and hormones. Upon activation, receptor signalling converges onto four Gα protein subtypes to regulate intracellular responses. Therefore, variation in a single Gα protein gene can potentially impact the function of numerous receptors. In this work, we have performed a multidimensional study of rare disease mutations in Gαs, a prototypical Gα protein. By integrating data from 3D structures, GPCR / G protein functional pairings, transcriptomics, biophysics, and molecular dynamics with systems pharmacology modelling, our results reveal why mutations impairing receptor / Gαs coupling result in highly specific context-based signalling defects. Furthermore, we show that mutations leading to the same rare disease can alter different signal transduction steps, highlighting the importance of patient-specific treatment strategies. By closely dissecting G protein coupling, our study provides a blueprint to interrogate GPCR pathway signalling diversity in different (patho)physiological contexts.

## INTRODUCTION

G protein-coupled receptor (GPCR) signalling modulates key physiological processes such as vision, olfaction, neurotransmission and metabolism. In this signalling system, hundreds of receptors have diversified to detect a wide collection of structurally-diverse signals which they relay to four different G protein types to elicit an array of intracellular responses. Since the landmark publication of the structure of a prototypical ternary complex involving the β2 adrenergic receptor bound to a Gs heterotrimer in 2011 (*1*), the GPCR signalling field has greatly advanced on the concerted effort to understand receptor / G protein coupling from multiple dimensions. On the one hand, information on the structure of GPCRs in complex with diverse G proteins has grown dramatically(*2*). On the other, systematic analyses of the functional coupling preferences of hundreds of receptors towards different G protein types have started uncovering the pairing rules governing GPCR signalling(*3–5*). Importantly, parallel developments in RNA sequencing and proteomics now allow us to assess how different signalling partners combine in specific cells, tissues or organisms (*6*), thus giving rise to context-specific responses to the same stimulus (*7*). Taken together, all these insights have boosted our knowledge of GPCR function and offer the opportunity to address new holistic questions on receptor signalling such as: how does structural variation at the receptor or G protein level translate into phenotypic variability? how many receptor inputs converge into specific G proteins? and how can the cell or tissue-specific expression of different components of the GPCR machinery influence the detection of specific signals and determine their functional outputs?

In this work, we have leveraged on state-of-the-art multidimensional data on GPCR / G protein coupling to perform a systems pharmacology analysis of single residue mutations in the prototypical Gαs protein leading to rare disease. These mutations (L388R and E392K), which have been detected in patients diagnosed with the endocrine disease pseudohypoparathyroidism type I c (PHPIc) (*8*), provide a unique case study to address the aforementioned key questions on the GPCR signalling system for three main reasons: i) although the resulting Gαs mutants are capable of promoting downstream signalling when they are directly stimulated by cholera toxin, they display a loss of function with regards to their activation by GPCRs (*8*); ii) considering their location in the helix 5 (H5) of Gαs, a key region determining the coupling and selectivity of G proteins towards GPCRs (*9*), both mutations have been postulated to impair Gαs – receptor interaction; and iii) by being encoded in an imprinted locus, Gαs transcripts show a highly specific pattern of expression. This results in Gαs from the maternal allele, which carries the mutations found in patients, being the only protein version expressed in some regions of the kidney, the thyroid, or the ovary, thus resulting in tissue-specific disease manifestations (*10*).

Our structural and network biology analyses reveal that L388R and E392K mutations have the potential to structurally and functionally impact Gαs protein coupling to multiple receptors. Importantly, by exploiting transcriptomics data to systematically evaluate which of these receptors are expressed in proximal renal tubules, where the main rare disease phenotype occurs, our results explain why the function of particular GPCRs, such as the parathyroid hormone receptor 1 (PTH1R), is specifically compromised in patients. Using a detailed characterisation of both Gαs mutants at multiple signal transduction steps by state-of-the-art biophysical methods, we uncover mutation-specific functional impairment, which we further rationalise through systems pharmacology modelling and molecular dynamics studies. Importantly, our models also allow suggesting patient-tailored treatment strategies based on individual rare disease variants and predicting how tissues lacking imprinting could be affected by Gαs heterozygosity. In summary, this study exemplifies how obtaining a multidimensional view of GPCR signalling allows dissecting receptor / G protein signal transduction and unravelling the functional impact of structural variation in GPCR pathway components from a context-based perspective. This, in turn, can foster a patient-centric understanding of phenotypic variation and disease, and inspire new strategies for personalised treatment.

## RESULTS

### Structural, functional and tissue-centric Gs coupling analysis

To gauge the structural relevance of mutations L388R and E392K on Gαs, we first mapped these positions into an active Gs protein structure (*11*) (PDB ID 6NBF), confirming both of them are located in the H5 region of the Gαs subunit (Fig. 1A). We next performed an analysis of all structurally solved GPCR / Gαs complexes to comprehensively evaluate the contribution of these residues towards receptor / G protein interactions. To do so, we analysed contacts between Gαs and receptors from 135 experimentally solved crystal and cryogenic electron microscopy (cryo-EM) structures, including 44 unique GPCRs (see Materials and Methods, **Supplementary Data 1**). Assigning each receptor and G protein residue a GPCRdb(*12*) and Common G Protein (CGN)(*13*) numbering scheme identifier respectively, allowed us to obtain residue contact frequencies across all available structures (Fig. 1B). A general overview of these contact frequencies confirms previous observations on the overall importance of the Gαs H5 in establishing an active GPCR / G protein interface (Fig. 1B, left). By focusing on this specific region, we observe how disease-related residues L388 and E392, corresponding to CGN G.H5.20 and G.H5.24 (Fig. 1B, right), establish frequent and extensive contacts with receptors. In the case of L388^G.H5.20^, interaction occurs with transmembrane (TM) helices 3, 5 and 6 of GPCRs, with contacts between this residue and receptor positions 5.61 and 5.64 found in 102 and 96 out of the 135 analysed structures. E392^G.H5.24^ contacts are established with TM6 and TM7 of receptors, as well as their helix 8. This Gαs residue shows a high contact frequency with this last helix, with interactions with positions 8.47 and 8.48 found in 109 and 97 analysed structures. Taken together, this analysis highlights the potential generalised impact of mutations in these Gαs positions when it comes to coupling to an array of GPCRs.

**Fig. 1.**
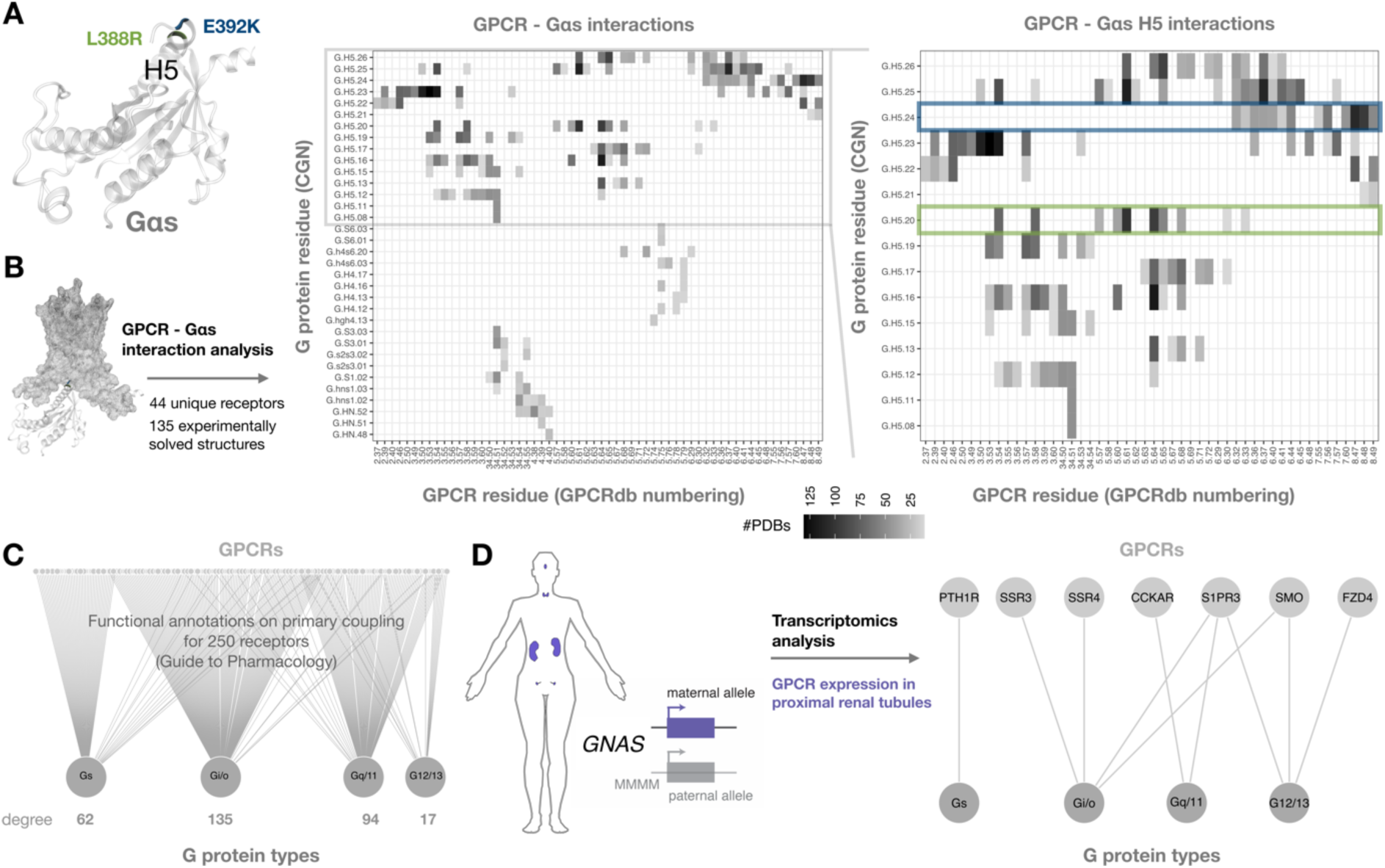
A context-based analysis of Gαs coupling. (**A**) Structural mapping of rare disease variants L388R (green) and E392K (blue) into an activated Gαs structure (white cartoon) (PDB ID 6NBF) showing both of them are located in the helix 5 (H5) of the Gαs subunit. (**B**) Interaction analysis of 135 experimentally solved GPCR - Gαs structures. Interactions present in at least 10 independent structures are shown in two heatmaps with residues annotated using GPCRdb generic numbering for receptors and the G protein common numbering (CGN) scheme for Gαs. Heatmaps include all interactions (left) and specific interactions between Gαs H5 and the receptor (right) with L388^G.H5.20^ and E392^G.H5.24^ interactions highlighted in green and blue boxes respectively. (**C**) Bipartite graph of primary couplings between GPCRs and G protein types as annotated in the Guide to Pharmacology. Degree values indicate the number of GPCRs that primarily couple to a particular G protein type. (**D**) Schematic representation of tissues displaying maternal monoallelic expression from the *GNAS* locus due to imprinting (left) and filtered bipartite graph of primary couplings between GPCRs and G protein types for receptors found to be expressed in proximal renal tubules via transcriptomics analyses (right).

To further assess the extent of this impact, we performed a systematic analysis of GPCR signalling preferences via the four different G protein types. We retrieved information on primary receptor couplings available via the GproteinDb (*14*) and originally annotated in the Guide to Pharmacology (*15*) (see Materials and Methods for further details) to generate a GPCR / G protein functional coupling graph (Fig. 1C). This revealed that the function of at least 62 different GPCRs, which preferentially couple to Gαs, could be theoretically affected by the rare disease mutations. We know, however, that considering tissue-specific context is highly relevant in the rare disease due to the fact that the *GNAS* locus is imprinted in a number of tissues leading to the expression of a single maternally-inherited Gαs version which is mutated in patients(*8*). This is particularly relevant in the proximal renal tubules of the kidney, where the disease results in disrupted calcium resorption. For this reason, we mined publicly available transcriptomics data from different human kidney segments (*16*) to filter the initial GPCR / G protein network so that it only included those receptors expressed in the proximal renal tubules (Fig. 1D). Remarkably, this revealed that the only receptor displaying primary Gαs coupling in this tissue was the parathyroid hormone receptor 1 (PTH1R), thus explaining previous observations on the unique role of this GPCR in the aetiology of the rare disease.

### Rare disease variants alter diverse signal transduction steps

Initial work characterising L388R and E392K mutations concluded these rare disease variants were likely to disrupt the receptor signalling process by preventing GPCR / G protein interaction(*8*) (Fig. 2A, step 1). To dissect their influence on PTH1R signal transduction processes, we first took advantage of mini-G proteins - engineered variants of the GTPase domains of Gα subunits (*17*) - to directly monitor Gαs recruitment to PTH1R excluding any interference from receptor coupling to additional G proteins. Through the use of mini-G proteins bearing both mutations, we were able to measure their interaction with PTH1R, as well as their recruitment to the plasma membrane, upon activation by parathyroid hormone 1-34 (PTH(1–34)) via bioluminescence resonance energy transfer (BRET) (see Materials and Methods). As expected, the L388R mutant showed impaired receptor recruitment upon agonist stimulation, which was also reflected by a diminished translocation to the plasma membrane marker, K-Ras (Fig. 2, B and C, green curve). Surprisingly, the E392K mutant showed a completely different behaviour, displaying higher receptor and plasma-membrane interaction levels than *wild type* mini-G (Fig. 2, B and C, blue curve).

**Fig. 2.**
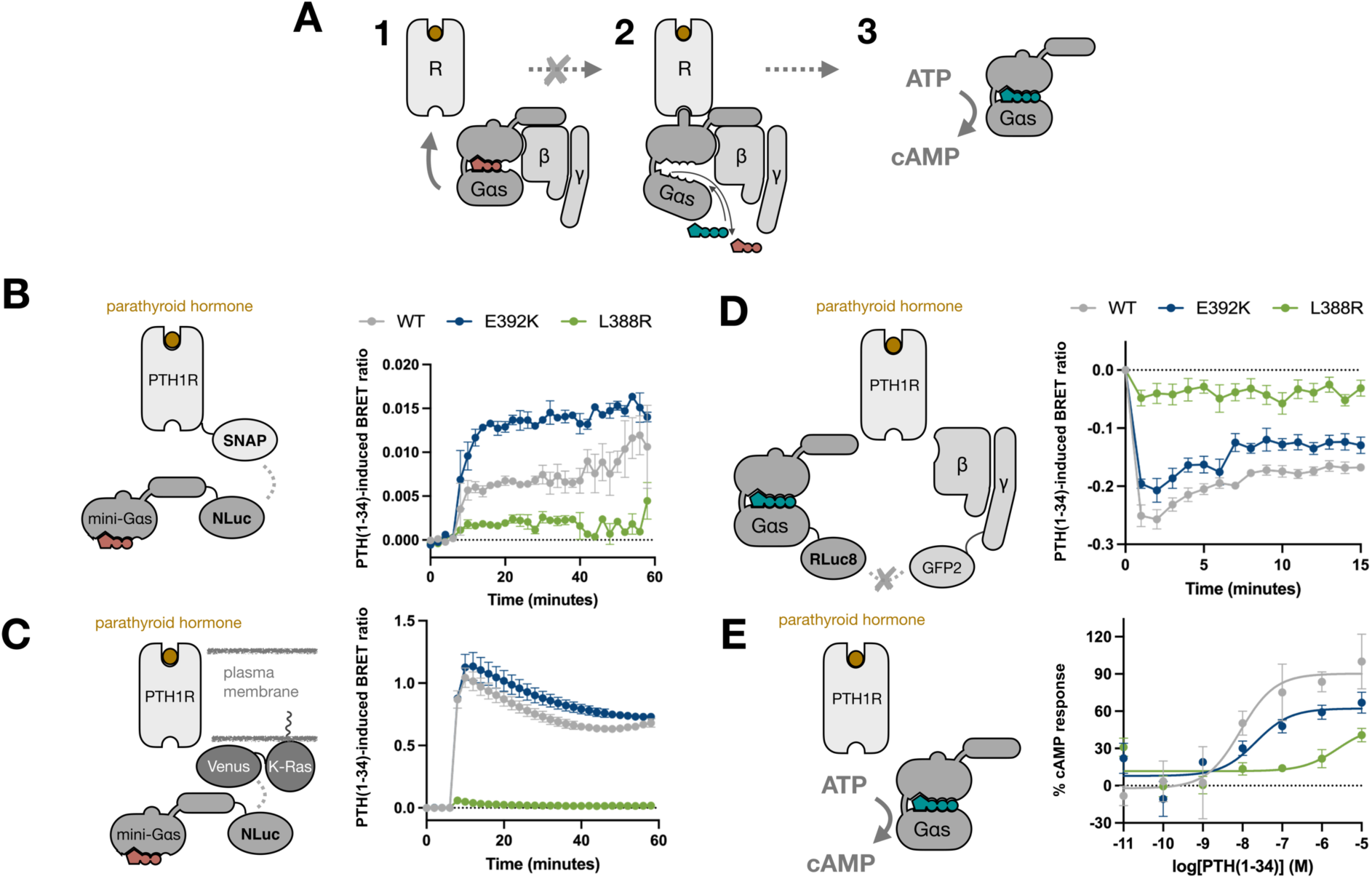
Monitoring differential receptor engagement in Gαs rare disease variants. (**A**) Schematic representation of key steps in the Gαs activation process including heterotrimeric G protein coupling to an activated receptor (R) (step 1), receptor-induced exchange of GDP (clay) for GTP (teal) (step 2), and GαsGTP-mediated cAMP production after dissociation from the Gβγ subunits (step 3). The grey cross represents the activation step reported to be compromised upon L388R or E392K mutation. BRET assays used to measure mini-Gαs (WT, E392K and L388R) translocation to PTH1R (**B**) or plasma membrane marker K-Ras (**C**) following stimulation with 1μM PTH (1–34). The BRET1 ratio was calculated as donor/acceptor wavelength and baseline-corrected with the value obtained with vehicle control. (**D**) Dissociation of WT heterotrimeric Gαs and Gαs containing either the E392K or L388R mutations, as determined using the TRUPATH G protein activation assay following stimulation of the PTH1R using 1μM PTH(1-34). The BRET2 ratio was calculated as donor/acceptor wavelength and baseline-corrected with the value obtained with DMSO control. (**E**) Differential cyclic adenosine monophosphate (cAMP) accumulation induced by WT Gαs or E392K and L388R mutants upon PTH1R stimulation. cAMP accumulation after 1hr stimulation with 10μM-0.01nM PTH(1-34) was measured using the LANCE cAMP kit in HEK293ΔGNAS cells transiently transfected with Gαs WT/E392K/L388R. The data was normalised to the maximum cAMP production observed for WT Gαs. All values are represented as mean ± SEM, with n=3 independent experimental repeats in duplicates.

To further investigate these differences, we monitored a subsequent step in receptor-mediated G protein activation by measuring the ligand-induced dissociation of the heterotrimeric Gs protein complex. To do so, we followed the loss of BRET signal between Gαs and Gβγ upon PTH1R stimulation with 1 μM PTH(1–34) using the TRUPATH platform (*18*). Consistent with other examples using TRUPATH to study Gαs activation, following stimulation with PTH(1–34), the *wild type* G protein displayed a rapid loss of BRET signal, indicative of dissociation between Gαs and Gβγ followed by a gradual rise to steady state. In line with our observations showing impaired L388R interaction with PTH1R, a Gαs version including this mutation displayed minimal ligand-induced G protein dissociation (Fig. 2D). The E392K mutant, conversely, showed an intermediate response, with ligand-induced Gαs / Gβγ dissociation being significantly lower than in the *wild type* G protein (as measured by area under the curve (AUC) *p* = 0.0014; *n* = 6, Supplementary Materials, fig. S1A). These results are in accordance with previous downstream observations for these mutants (*8*), where E392K and L388R respectively display an intermediate and more pronounced loss of function with regards to cyclic adenosine monophosphate (cAMP) accumulation (Fig. 2E, 67% and 41% of the *wild type* at the highest PTH(1–34) concentration). Notably, these differences were also reproduced by using the TRUPATH platform to monitor isoprenaline-induced Gs protein activation at the prototypical β2-adrenoceptor (Supplementary Materials, fig. S1B). Overall, our detailed analyses highlight how distinct mutations leading to the same rare disease can do so by disrupting separate steps in the signal transduction process.

### A systems pharmacology model for impaired G protein function

The observed diversity in mutant behaviour prompted us to further explore which specific steps in the receptor-mediated G protein activation cycle were impacted by the different rare disease variants. To rationalise these observations, we generated a systems pharmacology model of G protein activation using ordinary differential equations (Fig. 3A and Supplementary Materials, Tables S1 and S2, see Materials and Methods). Initially, the model was fitted to the data points from the TRUPATH experiments for the *wild type* Gαs (Fig. 3B) using optimisation algorithms to minimise the squared error between simulation and experimental data as described previously (*19*). Next, we systematically explored perturbations in model parameters that could explain the G protein activation differences measured in the two Gαs mutants (see Materials and Methods). Based on the data shown in Fig. 2B, we inferred that, for the L388R mutant, G protein binding to the agonist-stimulated GPCR was impaired. This behaviour was reproduced in our model by reducing k_G+_, the constant governing G protein binding to different receptor activation states, by 1000-fold (k_G+_ of 1.604×10^7^ M^-1^s^-1^ for the *wild type* G protein vs. 1.604 x10^4^ M^-1^s^-1^ for L388R) (Fig. 3C and Supplementary Materials, Table S3). For the E392K mutant, the observed reduction of Gαs-GTP production was replicated by decreasing the parameter describing the capacity of the G protein to be activated by a GPCR (k_GDA_) by 49-fold compared to the *wild type* (k_GDA+_ of 2.690×10^7^ s^-1^ vs 5.487×10^5^ s^-1^) (Fig. 3D and Supplementary Materials, Table S3). These parameter changes allow our simulated curves to closely fit the area under the curve values from the original TRUPATH experiments (Fig. 3E).

**Fig. 3.**
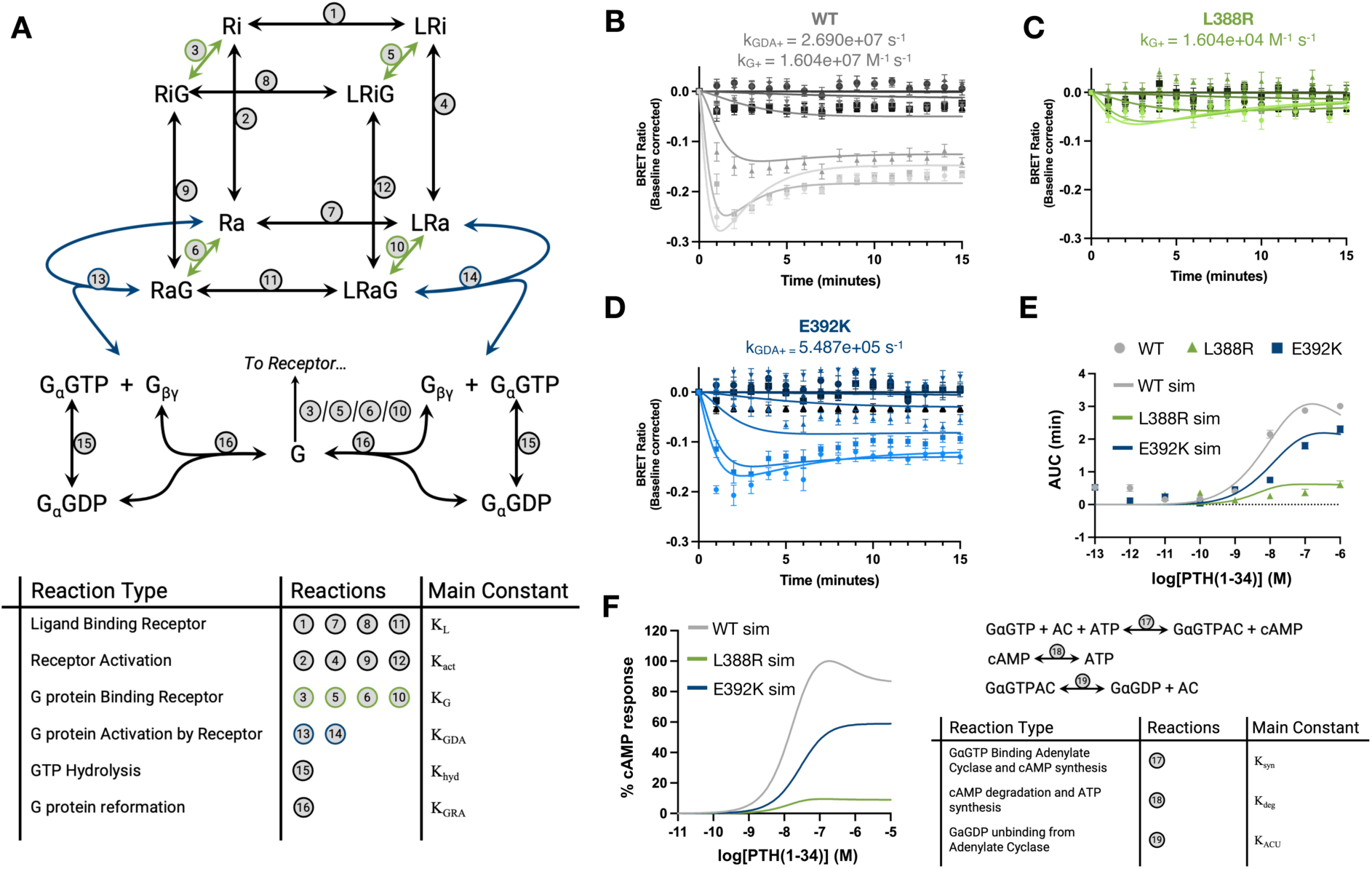
A systems pharmacology model of Gαs activation. (**A**) Cubic ternary complex model of GPCR signalling with extended reactions enabling simulation of the Gαs activation cycle (see Materials and Methods). Reactions are summarised in the table below with associated equilibrium constants. Reactions with shared parameters used to simulate all three variants are shown in grey. Reactions that have been changed to simulate variant effects are shown in green for L388R and in blue for E392K. (**B-D**) Time course values for the TRUPATH G protein activation assay following 15-minute stimulation of the PTH1R using 1μM – 0.1pM PTH(1-34) for WT (grey), L388R (green), E392K (blue) are represented as points. Solid lines have been fitted using the model shown in **a** with the parameter changes indicated in each panel. (**E**) Experimental dose response values for WT Gαs (grey) and Gαs containing L388R (green) and E392K (blue) mutations, measured as the area between the kinetic trace and baseline for each concentration (0.1pM-1μM of PTH(1-34)), are represented as points. Solid lines represent equivalent simulations for concentrations between 0.1pM and 1μM. (**F**) Simulated dose-response curves of cAMP production (measured at peak cAMP) after stimulation with 0.1pM to 1μM PTH(1-34). The right panel shows the additional simplified reactions (see Materials and Methods) appended to the model in (**A**) enabling simulation of cAMP production.

Furthermore, expanding our newly created model to estimate cAMP production levels by the *wild type* and mutated Gαs variants allowed us to reproduce our previously characterised signalling trends (Fig. 2E), which were also observed when the L388R and E392K mutations were originally characterised (*8*). In this way, this expanded model replicates the conservation of ligand-induced half maximal effective concentration (EC50) for the L388R and E392K mutants, together with the previously observed loss of function pattern for cAMP response at the highest PTH(1–34) concentration (Fig. 3F, 68% and 10% respectively compared to the *wild type*). Collectively, insights from our systems pharmacology modelling further illuminate how the L388R mutant could present a loss of function related to impaired receptor association, while the E392K mutation could lead to a suboptimal receptor-mediated activation of Gαs.

### Mutations alter GPCR – G protein interfaces and dynamics

To gain a structural understanding of the potential effects of our rare disease mutations on GPCR - Gαs interactions, we next took advantage of the structurally solved cryo-EM structure of PTH1R bound to the Gs protein and a PTH analogue (*11*) (PDB ID 6NBF). Modelling each mutation into the solved Gαs structure (see Materials and Methods) clearly suggested why one of the mutations may have a more pronounced impact on GPCR - Gαs complex formation: specifically, mutating a leucine into an arginine at position 388 of Gαs (G.H5.20 following the CGN), would replace a set of hydrophobic contacts with adjacent residues I320^3.58^ and L385^5.61^ in the receptor interface with a bigger, positively charged residue, thus generating a highly unfavourable interaction and potentially an intermolecular clash (Fig. 4A). Therefore, upon L388R mutation, GPCR - Gαs complexes could display big conformational rearrangements as compared to *wild type*. In contrast, although mutation from a glutamic acid into a lysine could modify interactions established by the *wild type* Gαs with the backbone of residues N463^8.47^ and G464^8.48^ in the receptor, this mutation could still be accommodated and allow for alternative interactions making it more complex to evaluate its functional impact by analysing the structural model.

**Fig. 4.**
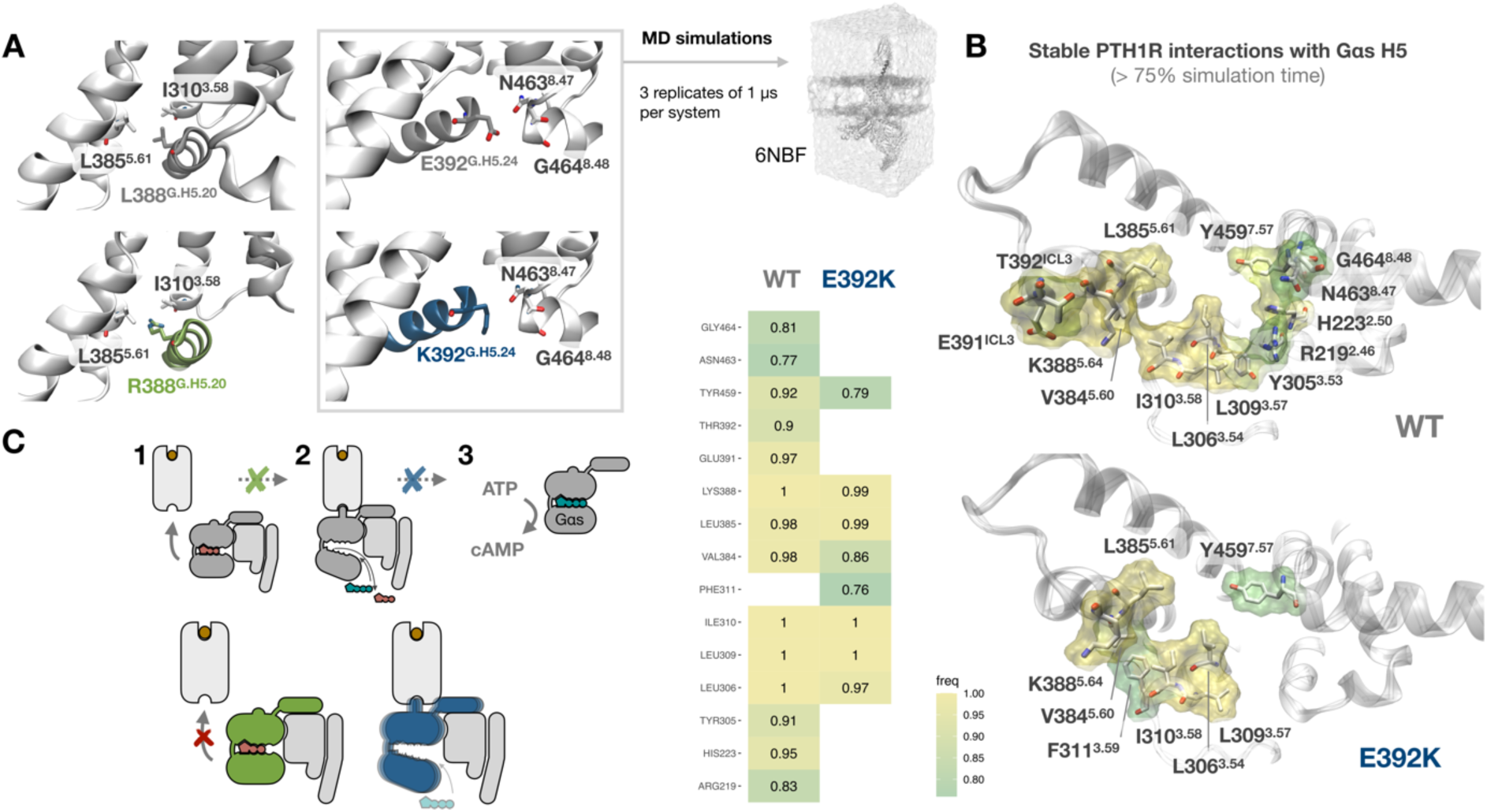
Structural analysis of PTH1R / Gαs interactions. (**A**) Structural model of mutation effects based on the structure of PTH1R bound to the Gs protein and a PTH analogue (PDB ID 6NBF). The top panels display L388 and E392 interactions between *wild type* Gαs protein (grey) and PTH1R receptor residues (white). The lower panels show a model of the potential effects of L388R (green) and E392K (blue) mutations on receptor interaction. Residues have been annotated using GPCRdb generic numbering for receptors and the G protein common numbering (CGN) scheme for Gαs. (**B**) Analysis of stable interactions of the PTH1R receptor with Gαs H5 during molecular dynamics (MD) simulations. Receptor residues establishing stable interactions (measured as residues that interact >75% of the simulation time across 3 independent MD replicates) are shown as sticks in an intracellular view of the 6NBF receptor structure and coloured according to interaction frequency (freq). The structural representation includes frequencies for the simulations including *wild type* (WT, top) and mutated (E392K, bottom) Gαs proteins. Specific frequency values for both systems are presented in the heatmap. (**C**) Schematic model of mutation-specific coupling deficiencies. While the L388R mutant (green) could present defective interactions with activated GPCRs (disrupting step 1 in the scheme), the E392K mutant (blue) could lead to impaired receptor-mediated Gαs protein activation (disrupting step 2).

The relatively subtle structural impact observed for the E392K mutation as compared to the *wild type* Gαs, prompted us to undertake a more detailed comparison of these two G proteins by assessing their stability in complex with PTH1R using classic molecular dynamics (MD) simulations (see Materials and Methods for a detailed protocol). Although the Gαs H5 maintained its helical conformation in both the *wild type* and the E392K mutant across 3 independent replicates of 1 μs (Supplementary Materials, fig. S2), analysis of stable contacts (occurring during >75% of the simulation time) between H5 and the receptor revealed a decrease in the number of contacts and in their frequency for the E392K mutant as compared to the *wild type* Gαs (Fig. 4B). Mapping these contact frequencies into the receptor intracellular interface shows how, although the E392K mutant preserves crucial stable interactions with residues in TM 3, 5 and 7; it lacks key contacts with TM2, the intracellular loop 3 (ICL3) and H8 that are established by the *wild type* Gαs. This incomplete engagement of H5 in the nucleotide-free state of the G protein suggests why receptor-mediated Gαs transition into a fully activated GTP-bound state is partially impaired upon E392K mutation. Altogether, our structural analyses are in line with our previous observations and support a mechanistic model in which one of the mutations, L388R, would result in Gαs losing its signalling capacity due to a lack of interaction with GPCRs (Fig. 4C, green), while the other, E392K, could be a product of GPCR – G protein complex instability resulting in a partial stalling of the Gα activation process (Fig. 4C, blue).

### Variant-specific drug response and cell signalling diversity

The differences in individual mutant behaviour observed in our multidimensional study open critical questions when it comes to suggesting new potential treatments with the ability to rescue Gαs signalling in disease-associated tissues. Even if both L388R and E392K mutations give rise to pseudohypoparathyroidism, successfully developing a potential ligand restoring cAMP production upon PTH1R stimulation could be highly mutation-dependent. To explore the ideal theoretical properties of such a ligand, we employed our previously developed systems pharmacology model to explore ligand parameter space (see Materials and Methods). Our simulations confirmed that altering ligand properties in the presence of the L388R mutation does not allow rescuing G protein activation; conversely, a compound promoting a slower deactivation of ligand-receptor complexes (corresponding to a 10.4-fold reduction in α_-_, the backwards cooperativity factor for ligand-bound receptor activation) could rescue Gαs activity in the E392K mutant (Fig. 5A). This predicted change in ligand properties is consistent with previously suggested therapeutic approaches based on the use of long-acting PTH derivatives to treat hypoparathyroidism patients (*20*). In more general terms, this exercise exemplifies why characterising specific causative mutations in individual patients can be critical when evaluating potential strategies for their treatment.

**Fig. 5.**
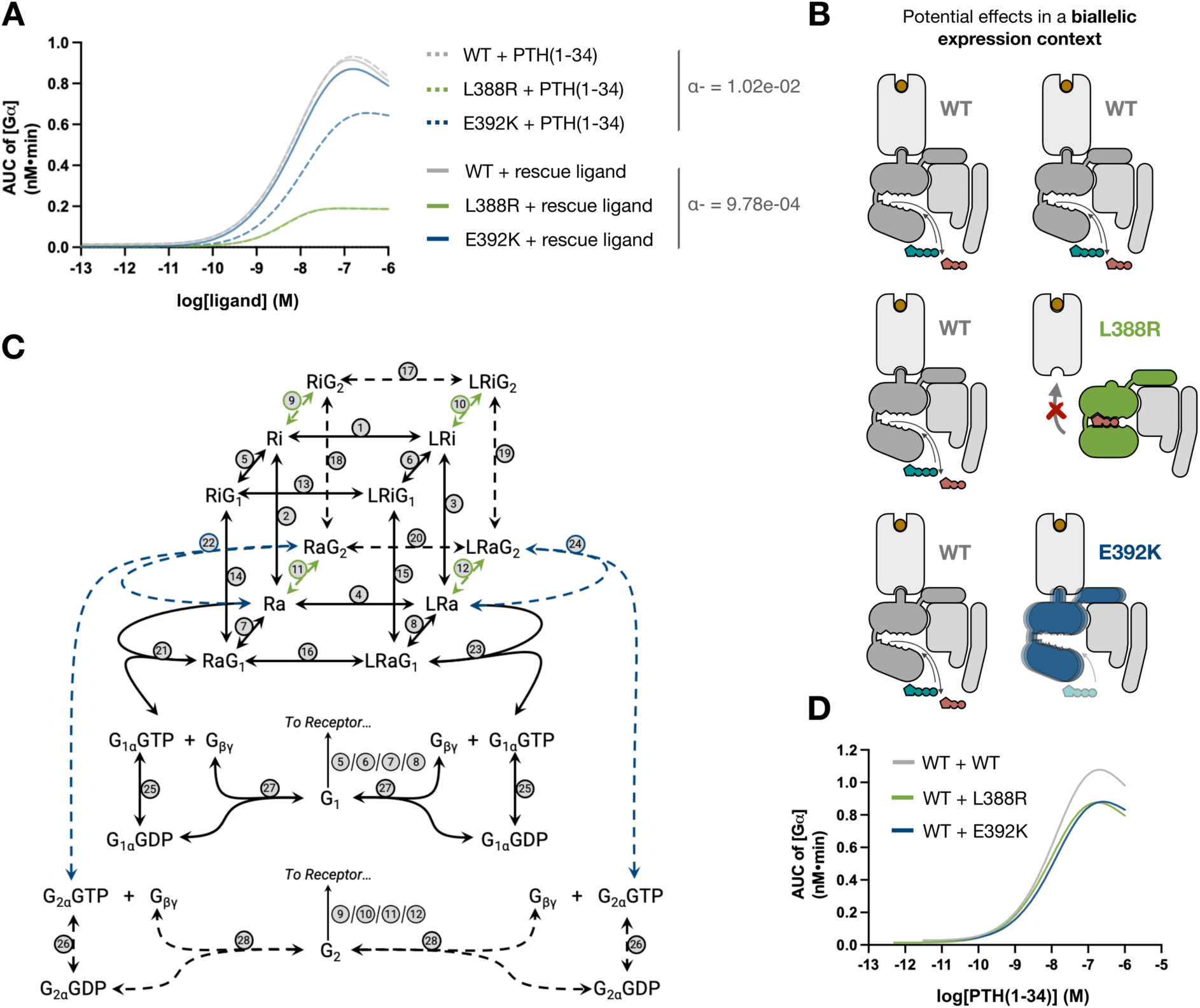
Modelling mutation rescue strategies and heterozygosity effects. (**A**) Simulation of ligand effects based on the model presented in Fig. 3. Original curves corresponding to total Gα accumulation in response to PTH(1-34) are shown as dashed lines, while signalling in response to a new theoretical rescue ligand fostering a slower deactivation of the ligand-receptor complex (∼10-fold decrease in the backwards cooperativity factor for ligand bound receptor activation (α_-_)) is shown as solid lines. (**B**) Schematic representation of expected Gαs protein diversity in healthy individuals (top) and rare disease patients bearing the L388R (middle, green) and E392K (bottom, blue) heterozygous mutations in tissues where the *GNAS* locus is not imprinted and there is biallelic Gαs expression. (**C**) Dual G protein cubic ternary complex model where the additional G protein can be a *wild type* (WT) Gαs or Gαs containing the L388R or E392K mutation. Reactions that have been changed to simulate variant effects are shown in green for L388R and in blue for E392K. (**D**) Simulated curves for total Gα production from the dual G protein model using the reactions presented in Fig. 3.

Another key consideration that can be explored with our models relates to the molecular phenotypes we can expect in non-imprinted tissues. In these tissues, patients will express two copies of Gαs, including a *wild type* version alongside the mutated one (Fig. 5B). To determine how the presence of two G protein versions with differing properties could alter overall signalling output, we built on our previous work simulating differential G protein coupling to the adenosine A1 receptor (*19*) to generate an extended model accounting for G protein heterozygosity (Fig. 5C and Supplementary Materials, Table S4, see Materials and Methods). Interestingly, this model shows that, in comparison to a system including two copies of the *wild type* Gαs, heterozygous expression of either *wild type* / L388R or *wild type* / E392K would produce an equivalent amount of total Gαs in response to PTH1R activation respectively, corresponding to 81,3% and 81,8% of the maximum response (Fig. 5D). In line with our previous mechanistic insights, this can be explained by a compensatory effect where the L388R mutant, which shows deficient GPCR binding, does not compete with the *wild type* G protein allowing it to freely engage with activated receptors; while the E392K mutant would still contribute to Gαs dissociation but with a lower efficacy than the *wild type*, thus interfering with the full activation of the pathway (Supplementary Materials, fig. S3). Notably, the fact that overall loss in G protein activation in this model (∼18%) is much lower than the one observed in the presence of individual Gαs mutants, can clarify why phenotypes associated with PHPIc mutations are predominantly constrained to imprinted tissues. These observations highlight the importance of accounting for heterozygosity when assessing variant effects present in the GPCR signalling machinery.

## DISCUSSION

In this work, we have combined comparative structural analyses, network biology reconstruction, transcriptomics mining, biophysical characterisation, systems pharmacology modelling and molecular dynamics simulations to explore how rare disease variation translates into context-specific signalling impairment in the GPCR system. In particular, our results reveal why specific receptors like PTH1R are uniquely affected by rare disease mutations in a central signalling node like the Gαs protein. We also show how L388R and E392K, two Gαs mutations giving rise to the same pathological phenotype, can do so by altering different steps of the signal transduction process. Furthermore, our models provide a mechanistic explanation for the alterations in molecular interaction properties and intermolecular complex stability that could respectively underlie defective signalling in each mutant. Importantly, our models also allow exploration of new theoretical strategies for mutation-centric signalling rescue and illustrate why monoallelic vs biallelic Gαs expression can result in tissue-specific dysfunction in rare disease patients.

Even if this study represents a paradigmatic example on the power of multidimensional analyses to understand receptor signalling variability, a deeper examination of downstream signalling pathways would be needed to fully characterise defective parathyroid hormone function in patients. This is due to the fact that additional signalling responses are mediated by PTH1R coupling to other intracellular partners like Gq/11 proteins and β-arrestins (*21*), whose interaction properties with the receptor could be indirectly modified by changes in Gαs engagement. Another key consideration is related to the ability of PTH1R to signal from intracellular compartments (*22*, *23*). Taking this into account, further dedicated work would be required to establish whether the changes we observe in the L388R and E392K Gαs mutants when it comes to PTH1R interaction could be compartment-specific. Additionally, the analysis pipeline presented in this work could be extended to other imprinted tissues to clarify additional observations of hormonal resistance found in some PHP patients (*10*). Finally, while our modelling and molecular dynamics simulation studies offer a mechanistic explanation for the observed effects of Gαs mutations on receptor signal transduction, additional structural evidence on G protein coupling intermediates could shed critical light into this process. This could be facilitated by new time-resolved cryo-EM strategies which have been recently applied to explore Gαs coupling to class A receptors (*24*).

Beyond our specific case study, this work can provide some fundamental guidance when it comes to characterising functional variation in the GPCR signalling system. This can be the case whether variation originates from disease mutations or from common polymorphisms present in the general population that could alter our predisposition to develop certain pathological phenotypes or to differentially respond to particular GPCR drugs in the clinic (*25*). First, our results exemplify the importance of monitoring orthogonal signalling outputs while characterising a set of variants related to a specific phenotype, as each of our biophysical analyses in isolation would have led us to contrasting conclusions when assessing the gain or loss of function of individual mutants. Second, our work highlights the need to consider the heterozygous expression of different versions of the same GPCR pathway component while attempting to determine whether variation will result in significantly different signalling effects. In this sense, the models presented here can serve as a blueprint to inspire future systems pharmacology analyses incorporating these considerations. Finally, our insights exemplify how identifying individual causal mutations in specific patients could be critical to guide the choice of therapeutic strategies and maximise their chances of success, supporting ongoing initiatives to transition into new personalised medicine treatment frameworks.

## MATERIALS AND METHODS

### Gαs - receptor protein contact analysis

Structures for 135 receptors in complex with Gαs (please see list in Data Availability) were downloaded from the RCSB PDB(*26*) (RCSB.org) and contacts between each receptor / Gαs protein pair were calculated using Arpeggio(*27*) with default parameters. Annotations regarding the G protein common numbering scheme were downloaded from the GproteinDb (*14*) resource and receptor residues were assigned generic numbering using the GPCRdb API (*28*). Our final analysis considered all receptor / Gαs protein interactions that occurred in at least 10 independent structures (*Supplementary Data 1*) and final interaction plots were obtained using RStudio 2022.07.2.

### Functional coupling and context-specific GPCR expression

Information on primary receptor couplings as annotated in the Guide to Pharmacology (*15*) was downloaded from GproteinDb (*14*) and represented as a bipartite graph using the igraph R library. Transcriptomics data from the human kidney was obtained via the Kidney Transcriptomics Data resource from the Epithelial Systems Biology Laboratory at NHLBI, NIH, Bethesda, MD (https://esbl.nhlbi.nih.gov/Databases/KSBP2/Targets/TranscriptomicData.html). To obtain a tissue-specific bipartite graph we used the dataset obtained by Cheval et al. (*16*) which is publicly available at https://doi.org/10.1371/journal.pone.0046876.s003 to search for GPCRs expressed in the S1 and S3 sub-segments of the proximal tubule.

### Mini-Gαs interactions with PTH1R and plasma membrane markers

A plasmid encoding PTH1R-SNAP was kindly provided by Ulrike Zabel and K-Ras-Venus was provided by Kevin Pfleger, University of Western Australia(*29*). NanoLuc (NLuc) luciferase-tagged mini-G probes were kindly provided by Nevin Lambert, Augusta University (*17*). The oligonucleotides for making the NLuc-mini-Gαs E392K and L388R mutations were designed using New England Biolabs’ NEBaseChanger online primer design tool, as forward *5’-CAGGCAGTATAAGCTGCTCTAAC-3’*, reverse *5’-GTAAGTCGCCTACGTAGA-3’* for NLuc-mini-Gαs E392K mutation and forward *5’-CGGATGCATCGCAGGCAGTAT-3’*, reverse *5’-GACAGCCCTGTAGTAAGTC-3’* for NLuc-mini-Gαs L388R mutation. Both mutations were generated by PCR mutagenesis and the sequences were confirmed by DNA sequencing at Source Biosciences (Cambridge, UK)

Human embryonic kidney (HEK) 293T cells were maintained in Dulbecco’s Modified Eagle’s Medium (DMEM) media (ThermoFisher, UK), supplemented with 10% Heat Inactivated Foetal Bovine Serum (FBS) (Sigma-Aldrich) and 100 U/ml penicillin and 0.1 mg/ml streptomycin (ThermoFisher, UK). Cells were maintained at 37 °C with 5% CO2, in a humidified atmosphere and passaged routinely. For BRET experiments, HEK293T cells were seeded at a density of 700,000 cell/well in a 6-well plate and grown overnight. The next day, cells were transfected with Lipofectamine 2000 (ThermoFisher) following the manufacturer’s protocol. For BRET experiments examining mini-G recruitment to PTH1R, 50ng of NLuc-tagged mini-G was transfected alongside 500ng C-terminally SNAP-tagged PTH1R, whereas, mini-G recruitment to plasma membrane, 50ng of NLuc-tagged mini-G was transfected alongside 500ng Venus-K-Ras and 300ng PTH1R. After 24 hours, cells were re-seeded onto white 96-well white polystyrene Nunc microplates (Sigma) precoated with poly-D-lysine (PDL)-coated, at a density of 100,000 cells/well in a complete FluoroBrite phenol red-free DMEM medium supplemented with 4 mM L-glutamine and 5% FBS medium and grown overnight. The next day, SNAP-tagged transfected cells were labelled with 1μM BG-Cy5 in complete FluoroBrite phenol red-free DMEM medium (without antibiotic) for 1 hour at 37 °C. Cells were then washed three times with complete FluoroBrite phenol red-free DMEM medium, followed by 80μL Hank’s balanced salt solution (HBSS) containing 10mM HEPES and 10μM furimazine/NanoGlo Luciferase Assay Substrate (Promega). BRET measurements were performed at 37 °C using a PHERAstar Microplate Reader (BMG Labtech) with a dual-luminescence readout BRET1 plus filter (460-490 nm band-pass, 520-550 nm long-pass). The BRET signal was recorded for 4 baseline measurements before the addition of 20μL PTH(1–34), and measured for an additional hour. The corresponding BRET ratio was calculated as the ratio of light emission from YFP (520 nm) over NLuc (460 nm). Net BRET ratio was baseline-corrected with vehicle-treated cells and normalized to the baseline values.

### Gαs activation analysis with TRUPATH

The pJG3.6-PTH1R construct was given to us by Dr. Simon Dowell (GSK, Stevenage, UK) and the β2-adrenoceptor (ADRB2) construct was gifted by Asuka Inoue (Tohoku University). The TRUPATH components including Gαs-Rluc8, Gβ3 and Gγ9-GFP2 were purchased as part of the TRUPATH biosensor kit from Addgene. The oligonucleotides for making the GαsE392K-Rluc8 and GαsL388R-Rluc8 mutations were designed using Agilent Technologies’ online primer design tool, as forward *5’-gagctattatagcagcttgtactgacgaaggtgca-3’*, reverse *5’-tgcaccttcgtcagtacaagctgctataatagctc-3’* for GαsE392K-Rluc8 and forward *5’-gctcgtactgacgacggtgcatgcgctga-3’* and reverse *5’-tcagcgcatgcaccgtcgtcagtacgagc-3’* for the GαsL388R-Rluc8 mutation. Both mutations were made using the QuikChange Lightening Site directed Mutagenesis Kit (Agilent Technologies) according to the manufacturer’s instructions and the sequences were confirmed by DNA sequencing at the Department of Biochemistry (University of Cambridge, UK). Peptide PTH(1–34) was purchased from Bachem (Bubendorf, Switzerland) and isoprenaline was purchased from Sigma (UK). Both were dissolved in dimethyl sulfoxide (DMSO) and stored as 1mM stock in -20°C.

Human embryonic kidney (HEK) 293T cells were maintained in Dulbecco’s Modified Eagle’s Medium (DMEM)/Hams F-12 nutrient mix (F12) GlutaMAXTM media (ThermoFisher, UK), supplemented with 10% Heat Inactivated Foetal Bovine Serum (FBS) (Sigma-Aldrich, Poole, Dorset, UK) and 1% antibiotic-antimycotic (AA) (Sigma, UK). Cells were maintained at 37 °C with 5% CO2, in a humidified atmosphere and passaged routinely. To perform the TRUPATH experiment, HEK293T cells were plated in a density of 1,500,000 cells/well in a 6-well plate and grown in complete DMEM /F-12 GlutaMAX^TM^ overnight. The seeded cells were transfected using 25 kDa polyethylenimine (PEI, Polysciences Inc., Germany) at a 6:1 ratio of PEI to DNA, diluted in 150mM NaCl. The receptor, Gαs-RLuc8 WT/E392K/L388R, Gβ3, Gγ9-GFP2 and pcDNA3.1 were transfected together with a ratio of 1:1:1:1:1 using 400 ng per construct. 24 hours post transfection, cells were trypsinised and re-seeded onto white 96-well plates (Greiner, UK) precoated with poly-L-lysine (PLL)-coated, at a density of 50,000 cells/well in a complete DMEM/F12 medium and grown overnight. On the next day, the cell culture media was removed, followed by wells washed with Hank’s balanced salt solution (HBSS). 80μl assay buffer (1× HBSS with calcium and magnesium, supplemented with 20 mM HEPES and 0.1% BSA with pH adjusted to 7.4) was added to each well, together with 10μl of coelenterazine 400a (Nanolight technology, USA) to a final concentration of 5 μM. For PTH1R experiments, the plates were then incubated in the dark for 5 minutes followed by the addition of 10 μl PTH(1–34) (ranging from 0.01nM to 10μM). For ADRB2 experiments, the TRUPATH G protein activation assay was performed using 10μM isoprenaline. The BRET signals were recorded every 60 seconds for 15 minutes on a Mithras LB940 plate reader and the corresponding BRET ratio was calculated as the ratio of light emission from GFP2 (515 nm) over Rluc8 (400 nm). Net BRET ratio was baseline-corrected with DMSO response and the negative peak in area under the curve (AUC) analysis was used to generate the dose-response curves. The dose-response curves were fitted with the three-parameter logistics equation built in Prism 9.3.1 (Graphpad Prism, San Diego, CA) for determining the response potency (pEC50). The statistical significance was calculated using ordinary one-way ANOVA with a Dunnett’s multiple comparisons test built in Prism 9.3.1.

### cAMP accumulation assay

The HEK293ΔGNAS cell line was kindly gifted by Asuka Inoue (Tohuku University). The pcDNA3.1-Gs-short construct was purchased from cDNA.org and the mutants GαsE392K and GαsL388R were generated and verified as described previously for the Gαs-RLuc8 mutants. HEK293ΔGNAS cells were maintained in DMEM/F12 GlutaMaxTM medium supplemented with 10% FBS and 1% AA. Cells were plated at a density of 300,000 cells/well in a 24-well plate and grown overnight prior to transfection. Transfection was performed with FugeneHD (Promega, UK) at a 3:1 ratio to DNA in accordance with the manufacturer’s instructions. PTH1R and GαsWT/GαsE392K/GαsL388R/pcDNA3.1 were transfected using 250ng per construct. After 48-hour transfection, cells were harvested and resuspended in the stimulation buffer (Phosphate buffer saline containing 0.1% BSA and 2.5mM isobutylmethylxanthine). The resuspended cells were then seeded at 1000 cells per well in 384-well white Optiplates (Perkin Elmer) and incubated with PTH(1–34) (ranging from 10μM to 10pM) or forskolin (Sigma, UK, ranging from 100μM to 100pM) for 1 hour at room temperature. The accumulated cAMP level was detected using the LANCE ultra cAMP detection kit on a Mithras LB 940 multimode microplate reader (Berthold Technologies). The response was normalised against the cAMP level produced from the stimulation of 100μM forskolin and fitted with the three-parameter logistics equation built in Prism 9.3.1 (Graphpad Prism, San Diego, CA).

### Systems pharmacology models of Gαs activation and signalling

We generated a mechanistic, kinetic model as presented in the reaction scheme in Fig. 3A using the reactions and parameters lists available in Tables S1 and S2. Using the law of mass action, we derived a system of Ordinary Differential Equations (ODEs) in COPASI 4.37 (Build 264) (*30*). The initial concentration of inactive receptor (Ri) and unbound heterotrimeric G protein (G) were both set to 415pM, as previously implemented (*19*). All receptor and G protein species were allowed to equilibrate for 10^6^ seconds before addition of ligand. The LSODA solver in COPASI(*30*) computationally solved our system of ODEs, yielding species’ concentrations over time.

To compare simulated G protein concentration with the TRUPATH experimental data, we applied the following linear transformation:

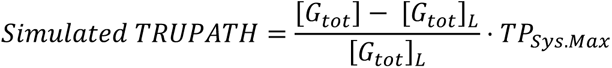

The concentration of total heterotrimeric G protein ([Gtot]) was normalised and baseline-corrected to the resting G protein concentration ([Gtot]L), where [Gtot]L is [Gtot] measured at the time of ligand addition. To obtain values directly comparable to the experimental data, we multiplied this normalised concentration by TPSys.Max, a constant representing the theoretical maximum TRUPATH output in our system if all G protein was activated. To fit simulated results to TRUPATH data on the *wild type* Gαs, we utilised COPASI’s parameter estimation task (*30*). Optimisation algorithms minimised the sum of squares (SoS) between experimental and simulated results at various ligand concentrations (1μM, 100nM, 10nM, 1nM). A final fit was found using a Genetic Algorithm with Stochastic Ranking with a SoS of 0.0733. To model experimental TRUPATH data for the L388R and E392K mutants, we scanned the parameter space for parameters representing properties upstream of G protein signalling: K_G_, β, γ, K_GDA_, and K_GRA_. (see Supplementary Materials, Table S2). Parameters of KG+ for the L388R mutant and KGDA+ for the E392K mutant (Supplementary Materials, Table S3) were selected based on agreement with mini-G experiments and close fitting to TRUPATH data, with a SoS between experimental and simulated data of 0.0956 for the L388R mutant and of 0.1041 for the E392K mutant.

To incorporate cAMP responses to the model, we appended simplified, collective reactions for cAMP synthesis (Fig. 3F, Supplementary Materials, Table S1). The initial concentration of AC was set to 4.15μM, while the initial concentration of ATP was set to 100μM. To examine the effects of theoretical rescue ligands in the model, we scanned model parameters representing properties of the ligand: KL, α and γ (Supplementary Materials, Table S2). Additionally, we derived a Dual G protein model from the original mechanistic model (Fig. 5C and Supplementary Materials, Table S4) by introducing an additional G protein at a concentration of 415pM. While Gα subunits of the two G proteins are distinct species, the Gβγ subunits from each G protein have identical properties. The final models have been made available in BioModels (*31*).

### Modelling and molecular dynamics of Gαs – PTH1R complexes

In order to assess the structural impact of the mutations on G protein / PTH1R interactions we first modelled both L388R and E392K mutations into the available cryo-EM structure of PTH1R bound to the Gs protein and a PTH analogue(*11*) (PDB ID 6NBF). These mutations were introduced using the Protein Builder function of the Chemical Computing Group’s Molecular Operating Environment (MOE) software (version 2022.02) and final structural representations were obtained with VMD1.9.4(*32*).

To set up the molecular dynamics simulation systems further protein curation was carried out using the MOE software including: (i) reverting A118 to the WT G188, (ii) modelling of missing sidechains and (iii) assignment of protonation states. Afterwards, internal water molecules were added using the HOMOLWAT server (*33*). Then, the curated protein was embedded into a solvated and ionized (150 mM NaCl) lipid bilayer composed of POPC (1-palmitoyl-2-oleoylphosphatidylcholine) using the CHARMM GUI membrane builder (*34*). The E392K mutant system was generated following the same protocol using the CHARMM GUI membrane builder. To run the molecular dynamics simulations, we employed the ACEMD v3.3 software (*35*). The systems underwent an initial energy minimization consisting of 500 steps to mitigate steric clashes and optimise the initial structure. Subsequently, an equilibration step was performed in the NPT ensemble for a duration of 10 nanoseconds. During this phase, restraints were applied to the protein Cα atoms at a force constant of 1 kcal/mol, as well as to the remaining heavy atoms at a force constant of 0.1 kcal/mol. These restraints were gradually reduced until reaching 0 kcal/mol over the 10 nanoseconds. For the production phase, the systems were allowed to freely evolve for a total time of 1 microsecond in three replicates. The production runs were carried out using a timestep of 4 femtoseconds in the NVT ensemble, where the temperature was kept constant at 300 K. Simulation data has been deposited at the GPCRmd repository (*36*).

To compare the stability of interactions between PTH1R and the H5 of Gαs we analysed each simulation frame across the 3 independent *wild type* and E392K mutant MD replicates with VMD1.9.4 (*32*) to select all receptor residues found within 4Å of residues 382 to 392 in the Gα protein. We considered an interaction to pass our stability filter when it occurred in more than 75% of the analysed frames. Per-residue secondary structure calculations over the simulation trajectories for the H5 of Gαs in *wild type* and E392K mutant conditions were obtained via the Timeline plugin implemented in VMD1.9.4. Final heatmap plots of those receptor residues establishing stable interactions with *wild type* and mutated Gαs were generated using RStudio 2022.07.2 and structural representations were obtained with VMD1.9.4 (*32*).

## ACKNOWLEDGEMENTS

The authors would like to thank Dr Maria Krantz for initial discussions on systems pharmacology models.

## Funding

Research in the laboratory of DC is supported by a Wellcome Trust Senior Research Fellowship (212313/Z/18/Z). XH is funded by a China Scholarship Council Cambridge International Scholarship. BM-L acknowledges financial support from an Industrial PhD Fellowship from the Spanish Ministry of Science and Innovation (DIN2021-011749) between the Research Program on Biomedical Informatics (GRIB) and Phamacelera S.L. JS acknowledges financial support from the Instituto de Salud Carlos III (ISCIII) (AC18/00030), and the Instituto de Salud Carlos III (ISCIII) co-funded by the European Union (PI18/00094). We gratefully acknowledge the support of the UK Biotechnology and Biological Sciences Research Council (BBSRC) (BB/W1014831/1 to GL, and a BBSRC-iCase studentship co-funded by AstraZeneca (V509334/1) to TR-N. GL is a Royal Society Industry Fellow (INF\R2\212001). MM-S is supported by a Royal Society University Research Fellowship (URF\R1\221205). She also acknowledges support from the Wellcome Trust Institutional Strategic Support Fund and the Isaac Newton Trust [22.23(d)]. This collaborative project was supported by the COST action CA 18133 ERNEST.

## Author contributions

MM-S designed and coordinated the research study; MM-S performed comparative structural biology, functional coupling, and transcriptomics analyses; SLO and DC designed and analysed mini-G experiments; SLO performed mini-G experiments; XH and GL designed and analysed TRUPATH and cAMP experiments; XH performed TRUPATH and cAMP experiments; TR-N and GL conceived and interpreted the systems pharmacology models; TR-N generated the systems pharmacology models; JS and MM-S designed and analysed the molecular dynamics simulations; JS and BM-L performed the molecular dynamics simulations; MM-S wrote the first version of the manuscript; all authors contributed to subsequent manuscript versions.

## Competing interests

Authors declare that they have no competing interests.

## Supplementary Materials

**Fig. S1.**
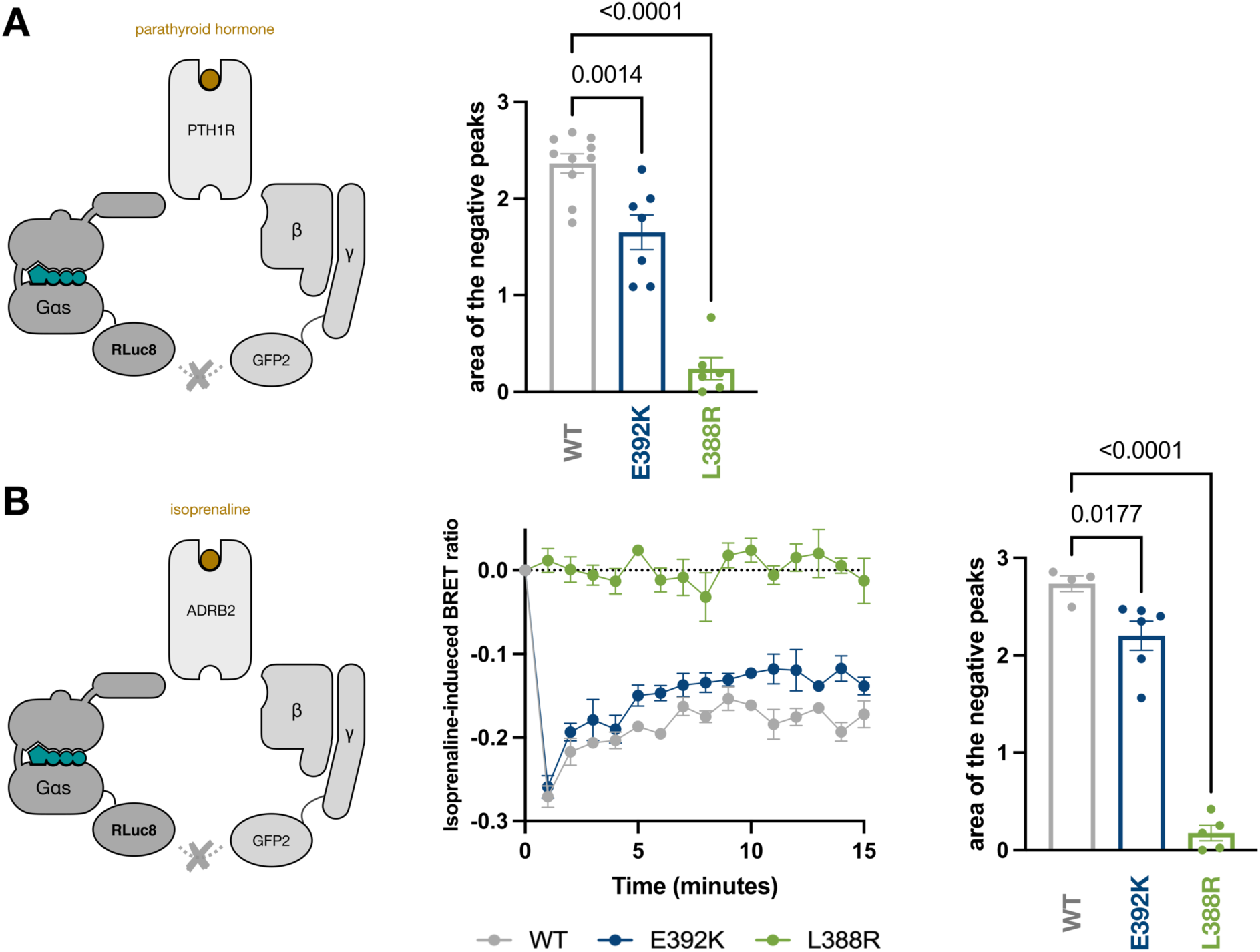
Activation of Gα variants by different Gs-coupled receptors. (**A**) Schematic representation of PTH1R-induced G protein activation measured in the TRUPATH assay and area of the negative peaks of 15min baseline-corrected BRET2 response induced by 1μM PTH(1–34) for the *wild type* (WT, grey), L388R (green) and E392K (blue) Gαs. (**B**) Schematic representation of β2 adrenergic receptor (ADRB2)-induced G protein activation measured in the TRUPATH assay. The central panel shows dissociation of WT heterotrimeric Gα_s_ (grey) and Gα_s_ containing either the L388R (green) and E392K (blue) mutations, as determined using the TRUPATH G protein activation assay following stimulation of ADRB2 using 10μM isoprenaline. The BRET2 ratio was calculated as GFP2/RLuc8 and baseline-corrected with the value obtained with DMSO as control. The right panel shows the area of the negative peaks of 15min baseline-corrected BRET2 response as in (**A**). for the *wild type* (WT, grey), L388R (green) and E392K (blue) Gαs. All values are represented as mean ± SEM, with n=3 independent experimental repeats conducted in duplicates. One-way AVOVA with a Dunnett’s multiple comparisons test was performed to compare the statistical significance of the differences in G protein activation level between Gαs WT and Gαs E392K/L388R.

**Fig. S2.**
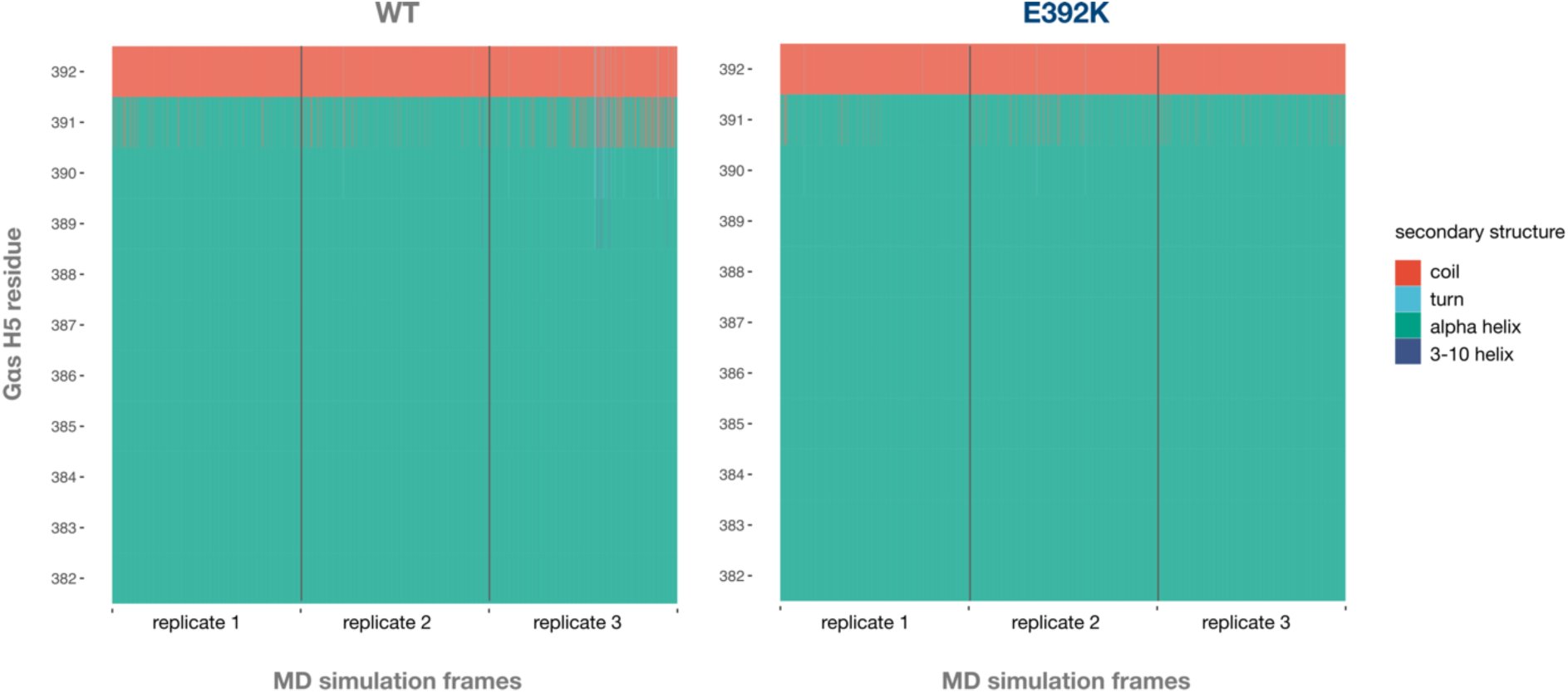
Gαs H5 secondary structure across MD simulation trajectories. Heatmap representation of residue-specific secondary structure assignments per simulation frame across 3 independent 1 μs replicates. Values were calculated with the VMD1.9.4 Timeline plugin and correspond to residues 382 to 392 of the helix 5 (H5) of Gαs from simulations with a *wild type* (WT, grey) and a mutated (E392K, blue) G protein.

**Fig. S3.**
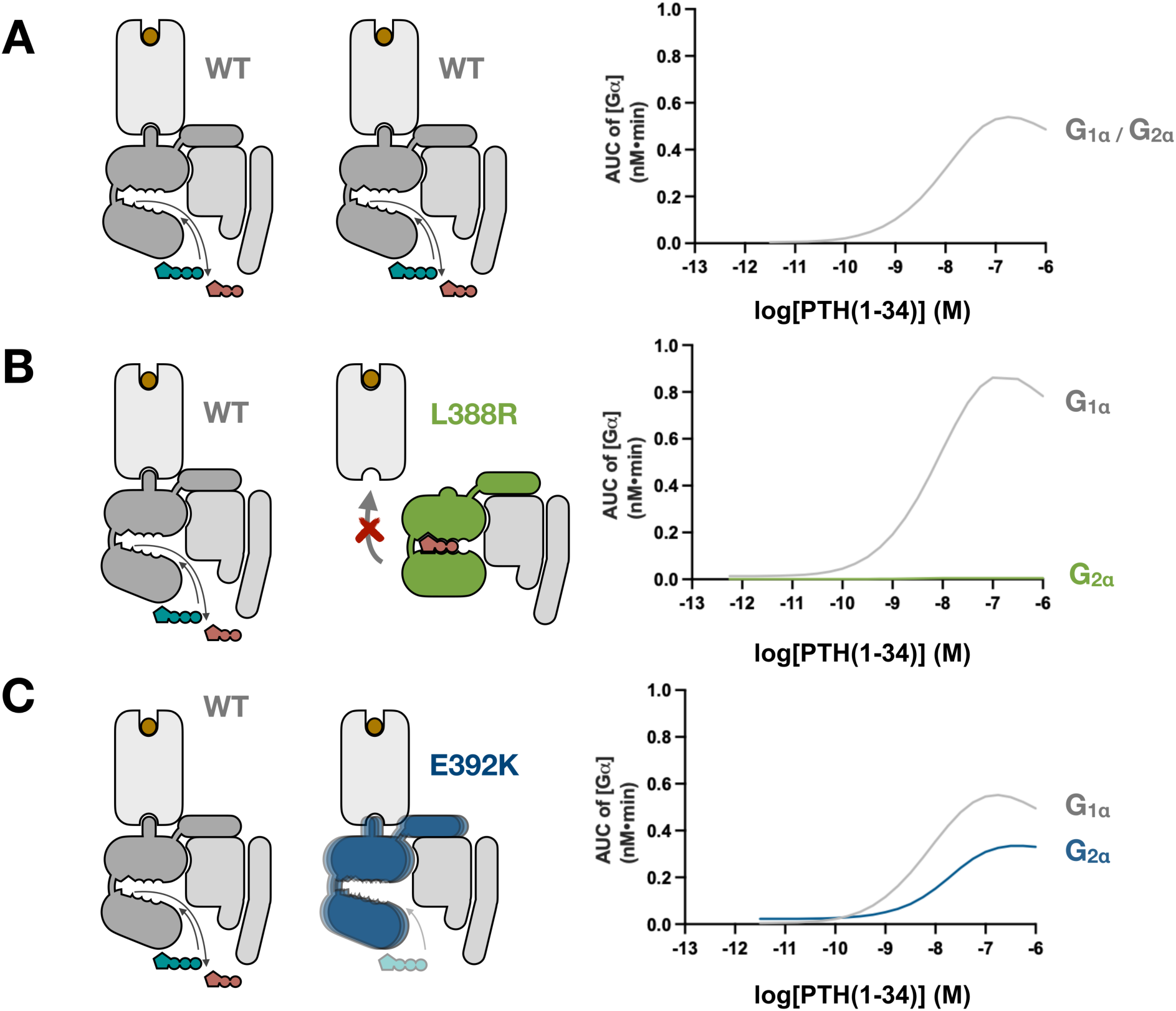
Activation of individual G proteins in the dual G protein model. Simulated curves for total Gα accumulation corresponding to G_1_ and G_2_ proteins in the dual G protein model presented in Fig. 5 (see Materials Methods). (**A**) Total Gα accumulation in the presence of two *wild type* (WT, grey) G proteins. (**B**) Total Gα accumulation in the presence of one *wild type* (WT, grey) G protein and a G protein containing the L388R mutant Gαs (green). (**C**) Total Gα accumulation in the presence of one *wild type* (WT, grey) G protein and a G protein containing the E392K mutant Gαs (blue).

**Table S1.** Reaction list for the G protein activation model.

| Reaction | Constituent Parameters | Value | Units |
| --- | --- | --- | --- |
| 1 $L + Ri \rightleftharpoons LRa$ | $k_{L+}$<br>$k_{L-}$ | 1.36E+07<br>3.61E+00 | $M^{-1} s^{-1}$<br>$s^{-1}$ |
| 2 $Ri \rightleftharpoons Ra$ | $k_{act+}$<br>$k_{act-}$ | 3.84E-06<br>2.54E+01 | $s^{-1}$<br>$s^{-1}$ |
| 3 $Ri + G \rightleftharpoons RiG$ | $k_{G+}$<br>$k_{G-}$ | 1.60E+07<br>5.33E-05 | $M^{-1} s^{-1}$<br>$s^{-1}$ |
| 4 $LRi \rightleftharpoons LRa$ | $k_{act+} \cdot \alpha_+$<br>$k_{act-} \cdot \alpha_-$ | 9.92E-03<br>2.59E-01 | $s^{-1}$<br>$s^{-1}$ |
| 5 $LRi + G \rightleftharpoons LRiG$ | $k_{G+} \cdot \gamma_+$<br>$k_{G-} \cdot \gamma_-$ | 1.58E+06<br>7.17E-07 | $M^{-1} s^{-1}$<br>$s^{-1}$ |
| 6 $Ra + G \rightleftharpoons RaG$ | $k_{G+} \cdot \beta_+$<br>$k_{G-} \cdot \beta_-$ | 1.26E+07<br>2.79E+02 | $M^{-1} s^{-1}$<br>$s^{-1}$ |
| 7 $L + Ra \rightleftharpoons LRa$ | $k_{L+} \cdot \alpha_+$<br>$k_{L-} \cdot \alpha_-$ | 3.51E+10<br>3.68E-02 | $M^{-1} s^{-1}$<br>$s^{-1}$ |
| 8 $L + RiG \rightleftharpoons LRiG$ | $k_{L+} \cdot \gamma_+$<br>$k_{L-} \cdot \gamma_-$ | 1.34E+06<br>4.86E-02 | $M^{-1} s^{-1}$<br>$s^{-1}$ |
| 9 $RiG \rightleftharpoons RaG$ | $k_{act+} \cdot \beta_+$<br>$k_{act-} \cdot \beta_-$ | 3.02E-06<br>1.33E+08 | $s^{-1}$<br>$s^{-1}$ |
| 10 $LRa + G \rightleftharpoons LRaG$ | $k_{G+} \cdot \beta_+ \cdot \gamma_+ \cdot \delta_+$<br>$k_{G-} \cdot \beta_- \cdot \gamma_- \cdot \delta_-$ | 1.24E+06<br>3.76E+00 | $M^{-1} s^{-1}$<br>$s^{-1}$ |
| 11 $L + RaG \rightleftharpoons LRaG$ | $k_{L+} \cdot \alpha_+ \cdot \gamma_+ \cdot \delta_+$<br>$k_{L-} \cdot \alpha_- \cdot \gamma_- \cdot \delta_-$ | 3.46E+09<br>4.96E-04 | $M^{-1} s^{-1}$<br>$s^{-1}$ |
| 12 $LRiG \rightleftharpoons LRaG$ | $k_{act+} \cdot \alpha_+ \cdot \beta_+ \cdot \delta_+$<br>$k_{act-} \cdot \alpha_- \cdot \beta_- \cdot \delta_-$ | 7.79E-03<br>1.36E+06 | $s^{-1}$<br>$s^{-1}$ |
| 13 $RaG \rightleftharpoons Ra + G\alpha GTP + G\beta\gamma$ | $k_{GDA+}$<br>$k_{GDA-}$ | 2.69E+07<br>1.00E-10 | $s^{-1}$<br>$M^{-2} s^{-1}$ |
| 14 $LRaG \rightleftharpoons LRa + G\alpha GTP + G\beta\gamma$ | $k_{GDA+}$<br>$k_{GDA-}$ | 2.69E+07<br>1.00E-10 | $s^{-1}$<br>$M^{-2} s^{-1}$ |
| 15 $G\alpha GTP \rightleftharpoons G\alpha GDP$ | $k_{hyd+}$<br>$k_{hyd-}$ | 2.42E+08<br>1.00E-10 | $s^{-1}$<br>$s^{-1}$ |
| 16 $G\alpha GDP + G\beta\gamma \rightleftharpoons G$ | $k_{GRA+}$<br>$k_{GRA-}$ | 5.15E-01<br>2.00E-05 | $M^{-1} s^{-1}$<br>$s^{-1}$ |
| 17 $G\alpha GTP + AC + ATP \rightleftharpoons G\alpha GTPAC + cAMP$ | $k_{syn+}$<br>$k_{syn-}$ | 1.00E+05<br>1.00E-20 | $M^{-2} s^{-1}$<br>$M^{-1} s^{-1}$ |
| 18 $cAMP \rightleftharpoons ATP$ | $k_{deg+}$<br>$k_{deg-}$ | 1.00E-01<br>1.00E-20 | $s^{-1}$<br>$s^{-1}$ |
| 19 $G\alpha GTPAC \rightleftharpoons G\alpha GDP + AC$ | $k_{ACU+}$<br>$k_{ACU-}$ | 1.00E-03<br>1.00E-20 | $s^{-1}$<br>$M^{-1} s^{-1}$ |

**Table S2.** Description of reaction parameters for the G protein activation model.

|  | Description | Value | Units |
| --- | --- | --- | --- |
| | $k_{L+}$ Ligand binding rate | 1.36E+07 | $M^{-1} s^{-1}$ |
| | $k_{L-}$ Ligand unbinding rate | 3.61E+00 | $s^{-1}$ |
| | $k_{act+}$ Receptor activation rate | 3.84E-06 | $s^{-1}$ |
| | $k_{act-}$ Receptor deactivation rate | 2.54E+01 | $s^{-1}$ |
| | $k_{G+}$ G protein binding rate | 1.60E+07 | $M^{-1} s^{-1}$ |
| | $k_{G-}$ G protein unbinding rate | 5.33E-05 | $s^{-1}$ |
| | $\alpha_+$ Forward cooperativity factor for ligand bound receptor activation | 2.58E+03 | - |
| | $\alpha_-$ Backwards cooperativity factor for ligand bound receptor activation | 1.02E-02 | - |
| | $\beta_+$ Forward cooperativity factor for G protein-bound receptor activation | 7.85E-01 | - |
| | $\beta_-$ Backwards cooperativity factor for G protein-bound receptor activation | 5.24E+06 | - |
| | $\gamma_+$ Forward cooperativity factor for ligand binding a G protein-bound receptor | 9.86E-02 | - |
| | $\gamma_-$ Backwards cooperativity factor for ligand binding a G protein-bound receptor | 1.35E-02 | - |
| | $\delta_+$ Forward cooperativity factor for ligand bound, G protein-bound receptor activation | 1.00E+00 | - |
| | $\delta_-$ Backwards cooperativity factor for ligand bound, G protein-bound receptor activation | 1.00E+00 | - |
| | $k_{GDA+}$ G protein dissociation rate from active, G protein-bound receptor (possibly ligand-bound) | 2.69E+07 | $M^{-2} s^{-1}$ |
| | $k_{GDA-}$ Reformation of active, G protein-bound receptor (possibly ligand-bound) | 1.00E-10 | $s^{-1}$ |
| | $k_{GRA+}$ Heterotrimeric G protein reassociation rate | 2.42E+08 | $M^{-1} s^{-1}$ |
| | $k_{GRA-}$ G protein spontaneous dissociation rate | 1.00E-10 | $s^{-1}$ |
| | $k_{hyd+}$ Rate of hydrolysis of GαGTP | 5.15E-01 | $s^{-1}$ |
| | $k_{hyd-}$ Spontaneous exchange rate of GDP for GTP | 2.00E-05 | $s^{-1}$ |
| | $k_{syn+}$ Collective rate of GαGTP binding to AC and of cAMP synthesis | 1.00E+05 | $M^{-2} s^{-1}$ |
| | $k_{syn-}$ Collective reverse rate of GαGTP binding to AC and of cAMP synthesis | 1.00E-20 | $M^{-1} s^{-1}$ |
| | $k_{deg+}$ Collective rate of cAMP degradation | 1.00E-01 | $s^{-1}$ |
| | $k_{deg-}$ Collective reverse rate of cAMP degradation | 1.00E-20 | $s^{-1}$ |
| | $k_{ACU+}$ Collective rate of GαGTP unbinding from AC and GTP hydrolysis to GDP | 1.00E-03 | $s^{-1}$ |
| | $k_{ACU-}$ Collective reverse rate of GαGTP unbinding from AC and GTP hydrolysis to GDP | 1.00E-20 | $M^{-1} s^{-1}$ |

**Table S3.**
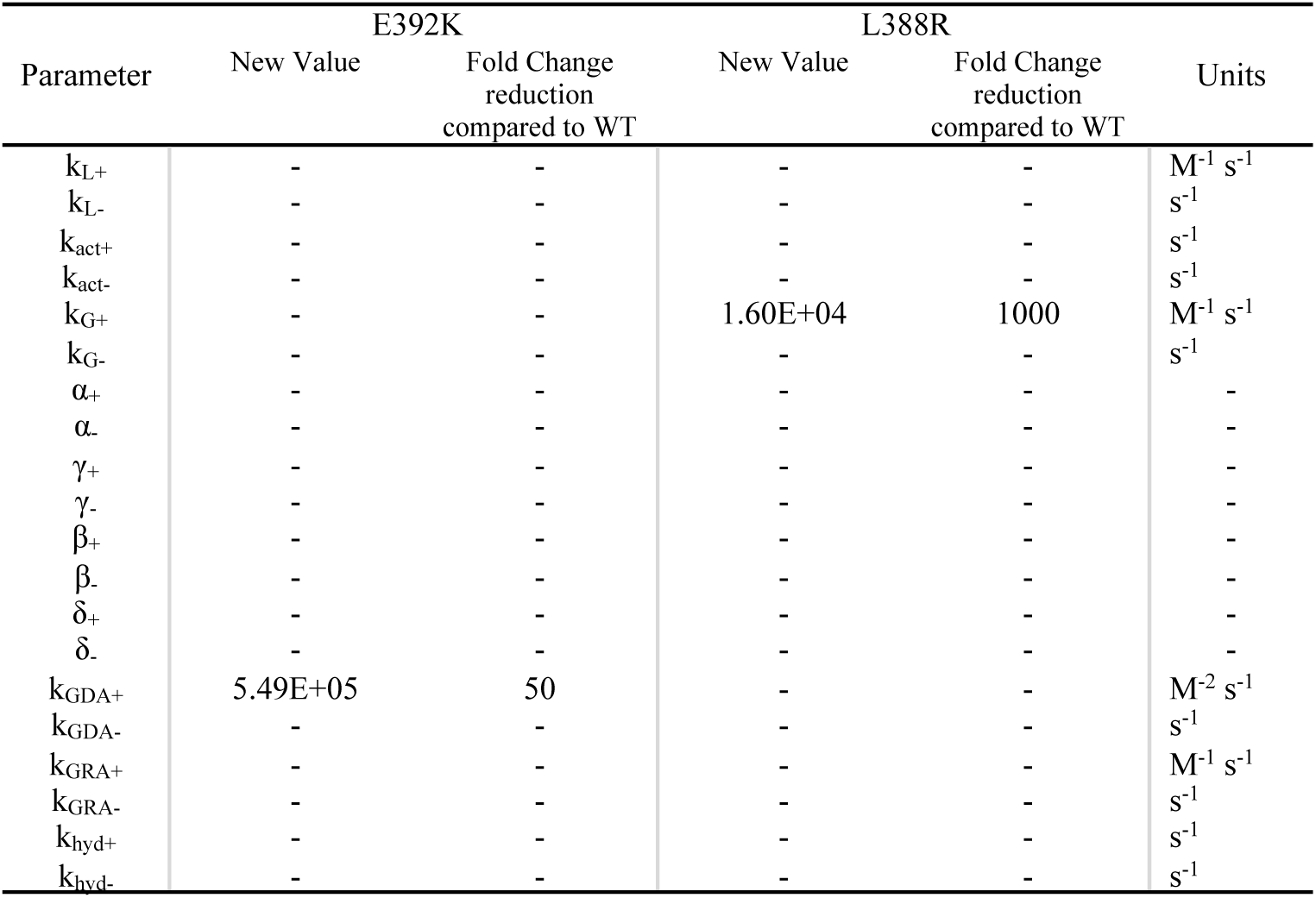
Parameter changes to simulate L388R and E392K mutants in the G protein activation model.

**Table S4.**
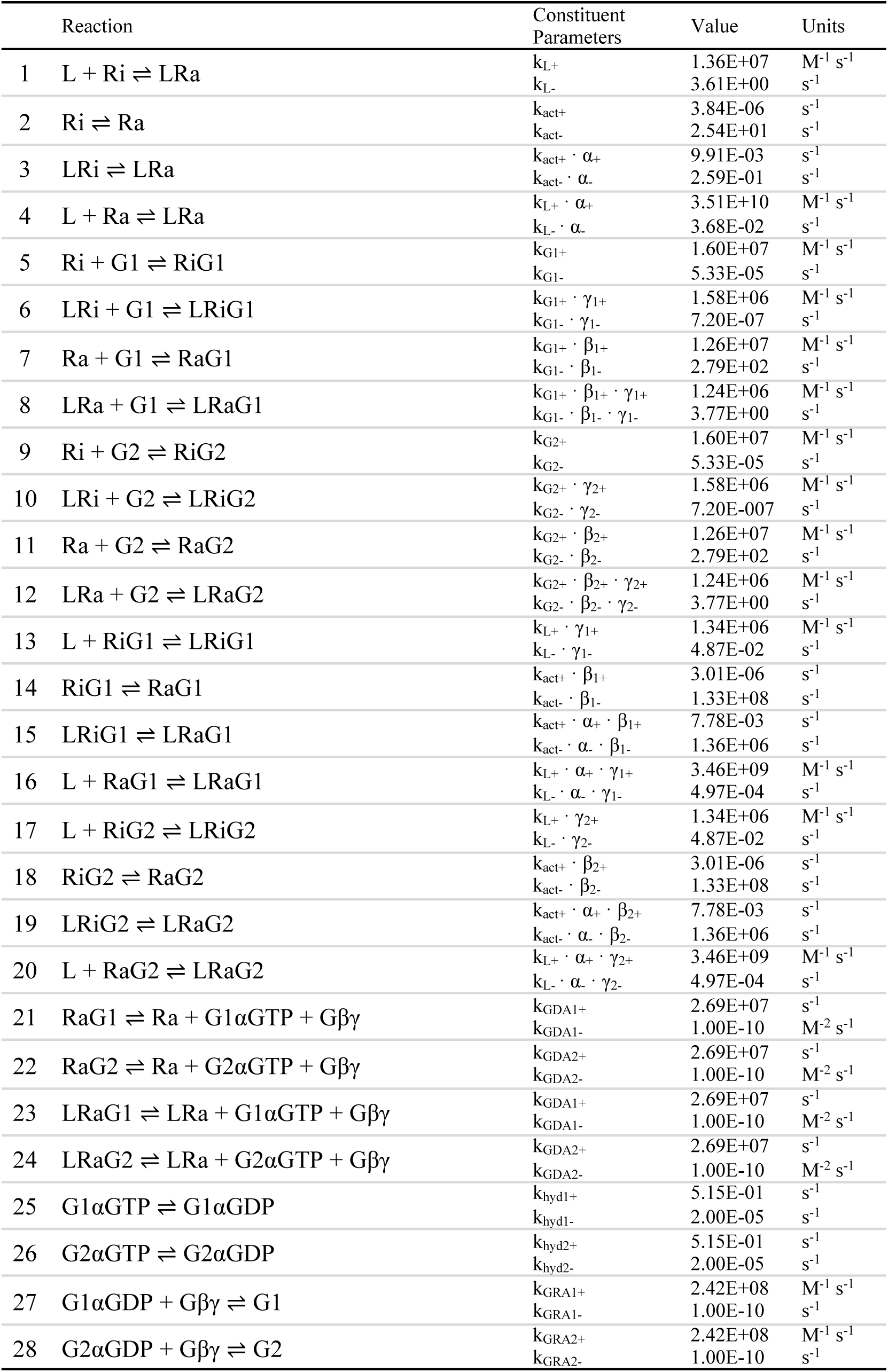
Reaction list for the dual G protein activation model.

| Reaction | Constituent Parameters | Value | Units |
| --- | --- | --- | --- |
| 1 $L + Ri \rightleftharpoons L Ra$ | $k_{L+}$<br>$k_{L-}$ | 1.36E+07<br>3.61E+00 | $M^{-1} s^{-1}$<br>$s^{-1}$ |
| 2 $Ri \rightleftharpoons Ra$ | $k_{act+}$<br>$k_{act-}$ | 3.84E-06<br>2.54E+01 | $s^{-1}$<br>$s^{-1}$ |
| 3 $LRi \rightleftharpoons L Ra$ | $k_{act+} \cdot \alpha_+$<br>$k_{act-} \cdot \alpha_-$ | 9.91E-03<br>2.59E-01 | $s^{-1}$<br>$s^{-1}$ |
| 4 $L + Ra \rightleftharpoons L Ra$ | $k_{L+} \cdot \alpha_+$<br>$k_{L-} \cdot \alpha_-$ | 3.51E+10<br>3.68E-02 | $M^{-1} s^{-1}$<br>$s^{-1}$ |
| 5 $Ri + G1 \rightleftharpoons RiG1$ | $k_{G1+}$<br>$k_{G1-}$ | 1.60E+07<br>5.33E-05 | $M^{-1} s^{-1}$<br>$s^{-1}$ |
| 6 $LRi + G1 \rightleftharpoons LRiG1$ | $k_{G1+} \cdot \gamma_{1+}$<br>$k_{G1-} \cdot \gamma_{1-}$ | 1.58E+06<br>7.20E-07 | $M^{-1} s^{-1}$<br>$s^{-1}$ |
| 7 $Ra + G1 \rightleftharpoons RaG1$ | $k_{G1+} \cdot \beta_{1+}$<br>$k_{G1-} \cdot \beta_{1-}$ | 1.26E+07<br>2.79E+02 | $M^{-1} s^{-1}$<br>$s^{-1}$ |
| 8 $L Ra + G1 \rightleftharpoons L RaG1$ | $k_{G1+} \cdot \beta_{1+} \cdot \gamma_{1+}$<br>$k_{G1-} \cdot \beta_{1-} \cdot \gamma_{1-}$ | 1.24E+06<br>3.77E+00 | $M^{-1} s^{-1}$<br>$s^{-1}$ |
| 9 $Ri + G2 \rightleftharpoons RiG2$ | $k_{G2+}$<br>$k_{G2-}$ | 1.60E+07<br>5.33E-05 | $M^{-1} s^{-1}$<br>$s^{-1}$ |
| 10 $LRi + G2 \rightleftharpoons LRiG2$ | $k_{G2+} \cdot \gamma_{2+}$<br>$k_{G2-} \cdot \gamma_{2-}$ | 1.58E+06<br>7.20E-007 | $M^{-1} s^{-1}$<br>$s^{-1}$ |
| 11 $Ra + G2 \rightleftharpoons RaG2$ | $k_{G2+} \cdot \beta_{2+}$<br>$k_{G2-} \cdot \beta_{2-}$ | 1.26E+07<br>2.79E+02 | $M^{-1} s^{-1}$<br>$s^{-1}$ |
| 12 $L Ra + G2 \rightleftharpoons L RaG2$ | $k_{G2+} \cdot \beta_{2+} \cdot \gamma_{2+}$<br>$k_{G2-} \cdot \beta_{2-} \cdot \gamma_{2-}$ | 1.24E+06<br>3.77E+00 | $M^{-1} s^{-1}$<br>$s^{-1}$ |
| 13 $L + RiG1 \rightleftharpoons LRiG1$ | $k_{L+} \cdot \gamma_{1+}$<br>$k_{L-} \cdot \gamma_{1-}$ | 1.34E+06<br>4.87E-02 | $M^{-1} s^{-1}$<br>$s^{-1}$ |
| 14 $RiG1 \rightleftharpoons RaG1$ | $k_{act+} \cdot \beta_{1+}$<br>$k_{act-} \cdot \beta_{1-}$ | 3.01E-06<br>1.33E+08 | $s^{-1}$<br>$s^{-1}$ |
| 15 $LRiG1 \rightleftharpoons L RaG1$ | $k_{act+} \cdot \alpha_+ \cdot \beta_{1+}$<br>$k_{act-} \cdot \alpha_- \cdot \beta_{1-}$ | 7.78E-03<br>1.36E+06 | $s^{-1}$<br>$s^{-1}$ |
| 16 $L + RaG1 \rightleftharpoons L RaG1$ | $k_{L+} \cdot \alpha_+ \cdot \gamma_{1+}$<br>$k_{L-} \cdot \alpha_- \cdot \gamma_{1-}$ | 3.46E+09<br>4.97E-04 | $M^{-1} s^{-1}$<br>$s^{-1}$ |
| 17 $L + RiG2 \rightleftharpoons LRiG2$ | $k_{L+} \cdot \gamma_{2+}$<br>$k_{L-} \cdot \gamma_{2-}$ | 1.34E+06<br>4.87E-02 | $M^{-1} s^{-1}$<br>$s^{-1}$ |
| 18 $RiG2 \rightleftharpoons RaG2$ | $k_{act+} \cdot \beta_{2+}$<br>$k_{act-} \cdot \beta_{2-}$ | 3.01E-06<br>1.33E+08 | $s^{-1}$<br>$s^{-1}$ |
| 19 $LRiG2 \rightleftharpoons L RaG2$ | $k_{act+} \cdot \alpha_+ \cdot \beta_{2+}$<br>$k_{act-} \cdot \alpha_- \cdot \beta_{2-}$ | 7.78E-03<br>1.36E+06 | $s^{-1}$<br>$s^{-1}$ |
| 20 $L + RaG2 \rightleftharpoons L RaG2$ | $k_{L+} \cdot \alpha_+ \cdot \gamma_{2+}$<br>$k_{L-} \cdot \alpha_- \cdot \gamma_{2-}$ | 3.46E+09<br>4.97E-04 | $M^{-1} s^{-1}$<br>$s^{-1}$ |
| 21 $RaG1 \rightleftharpoons Ra + G1\alpha GTP + G\beta\gamma$ | $k_{GDA1+}$<br>$k_{GDA1-}$ | 2.69E+07<br>1.00E-10 | $s^{-1}$<br>$M^{-2} s^{-1}$ |
| 22 $RaG2 \rightleftharpoons Ra + G2\alpha GTP + G\beta\gamma$ | $k_{GDA2+}$<br>$k_{GDA2-}$ | 2.69E+07<br>1.00E-10 | $s^{-1}$<br>$M^{-2} s^{-1}$ |
| 23 $L RaG1 \rightleftharpoons L Ra + G1\alpha GTP + G\beta\gamma$ | $k_{GDA1+}$<br>$k_{GDA1-}$ | 2.69E+07<br>1.00E-10 | $s^{-1}$<br>$M^{-2} s^{-1}$ |
| 24 $L RaG2 \rightleftharpoons L Ra + G2\alpha GTP + G\beta\gamma$ | $k_{GDA2+}$<br>$k_{GDA2-}$ | 2.69E+07<br>1.00E-10 | $s^{-1}$<br>$M^{-2} s^{-1}$ |
| 25 $G1\alpha GTP \rightleftharpoons G1\alpha GDP$ | $k_{hyd1+}$<br>$k_{hyd1-}$ | 5.15E-01<br>2.00E-05 | $s^{-1}$<br>$s^{-1}$ |
| 26 $G2\alpha GTP \rightleftharpoons G2\alpha GDP$ | $k_{hyd2+}$<br>$k_{hyd2-}$ | 5.15E-01<br>2.00E-05 | $s^{-1}$<br>$s^{-1}$ |
| 27 $G1\alpha GDP + G\beta\gamma \rightleftharpoons G1$ | $k_{GRA1+}$<br>$k_{GRA1-}$ | 2.42E+08<br>1.00E-10 | $M^{-1} s^{-1}$<br>$s^{-1}$ |
| 28 $G2\alpha GDP + G\beta\gamma \rightleftharpoons G2$ | $k_{GRA2+}$<br>$k_{GRA2-}$ | 2.42E+08<br>1.00E-10 | $M^{-1} s^{-1}$<br>$s^{-1}$ |

## Notes

### Competing Interest Statement

The authors have declared no competing interest.

